# Increasing flow rate reduces biofouling and colonization by filamentous bacteria in drippers fed with reclaimed wastewater

**DOI:** 10.1101/2020.06.02.130013

**Authors:** Kévin Lequette, Nassim Ait-Mouheb, Nathalie Wéry

## Abstract

The clogging of drippers due to the development of biofilms reduces the benefits and is an obstacle to the implementation of drip irrigation technology. The geometry of the dripper channel has an impact on the flow behaviours and head loss. The objective of this study was to analyse the influence of hydrodynamic parameters of three types of drippers (flow rates of 1, 2 and 4 l.h^-1^) fed by reclaimed wastewater on biofilm development kinetics and on the bacterial community. Using optical coherence tomography, we demonstrated that the inlet of the drippers (mainly the first baffle) and vortex zones are the most sensitive area for biofouling. Drippers with the lowest flow rate (1 l.h^-1^) and the smallest channel section were the favourable areas to biofouling. The low inlet velocity (0.34 m.s^-1^) in this type of dripper compared to 2 l.h^-1^ (0.61 m.s^-1^) and 4 l.h^-1^ (0.78 m.s^-1^) drippers can favour the deposition and development of biofilms. In addition, the water velocity influenced the structure of the bacterial communities in the biofilm. Low velocity (0.34 m.s^-1^) favoured the presence of *Hydrogenophaga* and *Pseudoxanthomonas* genera at the early stage of biofilm formation and filamentous bacteria belonging to Chloroflexi phylum at the end. So, maintaining a high flow rate and using drippers with a large flow cross-section is an effective way to control the development of biofilms by limiting the presence of filamentous bacteria.

## 1. Introduction

The scarcity of water resources is driving the development of water-saving irrigation techniques that enable integrated management of water resources. The reuse of reclaimed wastewater (RWW) is an appropriate way to alleviate the problem of scarce water resources especially in arid and semi-arid regions (Worako, 2015). Using RWW has several advantages including reducing pressure on freshwater resources and on the need for nutrients (e.g. nitrogen, phosphorus) for plant growth (Lazarova and Bahri, 2005) and can improve crop yields (Wang et al., 2013). Drip irrigation is the most water efficient (up to >90%) and safest technique for irrigation using RWW (Lamm et al., 2007). Drippers are usually composed of a labyrinth-channel and an outlet basin compartment. The geometry of the channel promotes the dissipation of the turbulent kinetic energy due to the zigzag flow path. However, the labyrinth channel is narrow (cross-section of around 1 mm) and the flow is thus sensitive to clogging (Liu and Huang, 2009; Niu et al., 2013), which reduces irrigation uniformity (Dosoretz et al., 2010) and requires more frequent maintenance (e.g. cleaning, replacing irrigation lines).

The process leading to clogging of the drippers is complex and the phenomenon is influenced by several parameters including water quality (concentration of suspended solids, chemical composition) (Lamm et al., 2007), the geometry of the dripper (Wei, 2011) as well as biological processes (Katz et al., 2014), the latter are the most difficult to control when RWW is used (Lamm et al., 2007). Previous studies have shown that flow velocity has an impact on the growth of biofilm in both drip-irrigation pipes (Li et al., 2012; Mahfoud et al., 2009) and in drippers (Gamri et al., 2014). The design of the geometry flow path influences the hydrodynamic flow and hence the development of biofilm (Bounoua et al., 2016). Computational studies of fluid dynamics have shown that the flow in the labyrinth-channel consisted of a main high velocity flow and vortex zones (Al-Muhammad et al., 2016) and that the development of biofilm is favoured in vortex areas due to reduced velocities and shear forces (Ait-Mouheb et al., 2018; Qian et al., 2017).

Flow behaviour not only influences the development of biofilms but also the microbial composition of the biofilms. Previous research on drinking water biofilms has shown that hydrodynamic conditions can drive the microbial community of the biofilms (Besemer, 2015; Rickard et al., 2004). However, in the context of the use of RWW, few authors have examined the influence of dripper parameters (i.e. flow rate, cross section, and geometry of the milli-channel) on microbial composition. Previous studies have described the microbial communities present in biofilms of different drippers by analysing phospholipid-derived fatty acids (PLFAs) (Yan et al., 2010, 2009; Zhou et al., 2017). These authors showed that three to seven types of PLFAs associated with Gram-negative bacteria were common in biofilms inside drippers, suggesting a significant role for these bacteria in biofilm development. However, in order to develop effective biofilm control strategies, deeper community characterization is required.

Recent study used high-throughput sequencing to describe the bacterial communities present in dripper biofilms to improve knowledge on clogging (Lequette et al., 2019; Song et al., 2019).

Optical methods (i.e. scanning electron microscopy) are commonly used to investigate the structure and the composition of biofilms at a local scale (Yan et al., 2009). But these studies underlined the importance of studying biofilm at mesoscale (millimetre range) in order to better understand the structure of biofilm, its function and the mechanisms behind its development (Morgenroth and Milferstedt, 2009). Recent studies used Optical coherence tomography (OCT) makes it possible to monitor biofilm formation non-invasively at a mesoscale and without staining (Derlon et al., 2012; Dreszer et al., 2014; Qian et al., 2017; West et al., 2016).

The objectives of the present study were to determine the effect of hydraulic parameters (1) on biofouling along the channel, (2) on the microbial communities in biofilms formed in irrigation systems fed by RWW. A non-destructive microscopic time-monitoring observation system was developed to study biofilms in commercial drippers with different flow rates (1, 2 and 4 l.h^-1^) with known cross-sections. The combined use of the OCT method and high-throughput sequencing made it possible to monitor the development of the biofilm under different hydraulic conditions while assessing the impact of these conditions on microbial composition.

## 2. Materials and Methods

### 2.1. Experimental setup

#### 2.1.1. Experimental setup and irrigation procedure

Non-pressure-compensating (NPC) drippers (model D2000, Rivulis Irrigation SAS, Lespinasse, France) with different flow rates (1, 2 and 4 l.h-1) were selected (Table 1). The development of biofilm in commercial drippers cannot be observed over time because the drippers are walled inside opaque irrigation pipes (black polyethylene tubes). The NPC drippers were therefore placed in a transparent tube (internal diameter 15mm, TubClair, Vitry-le-François, France), allowing us to see biofilm development along the channel over time. A solid tube with an internal diameter of 19mm (ABS+, Stratasys U-print SE plus, Stratasys Ltd, Eden Prairie, Minnesota, US) pressed the dripper against the walls of the transparent tube (Figure 1.A). The water flowed through the dripper grids connected to the milli-channel. Finally, an outlet hole (internal diameter 2mm) was pierced in the transparent tube above the outlet basin. Nine drippers were connected to each type of dripper using polyethylene tubing (length 10cm, internal diameter 20mm) and valves (internal diameter 20mm) (total length 3m). The valves were used to keep the drippers under water when they were disconnected from the irrigation line for analysis. Each of the three lines was connected to a separate tank (total volume 601) and a pump (JetInox 82M, DAB, Saint-Quentin-Fallavier, France) (Figure 1.B). A disk filter (mesh size 0.13mm) was installed to reduce the physical clogging of emitters following the technical recommendations of this type of dripper. The inlet pressure was set at 0.08 MPa with a pressure gauge. A gutter was installed below each lateral line to collect the water discharged from the drippers during the experiments. The gutters returned the discharged RWW to the respective tanks, enabling the recycling of water. The lines were supplied twice a day five days out of seven for 1h, with an interval of 6h off. The outlet flow rate from the drippers was measured weekly to evaluate emitter performance. Drippers are considered clogged when there is a decrease of at least 25% in the expected flow rate. The system was not flushed during the entire four-month experiment (from April to August 2018).

**Figure 1.**
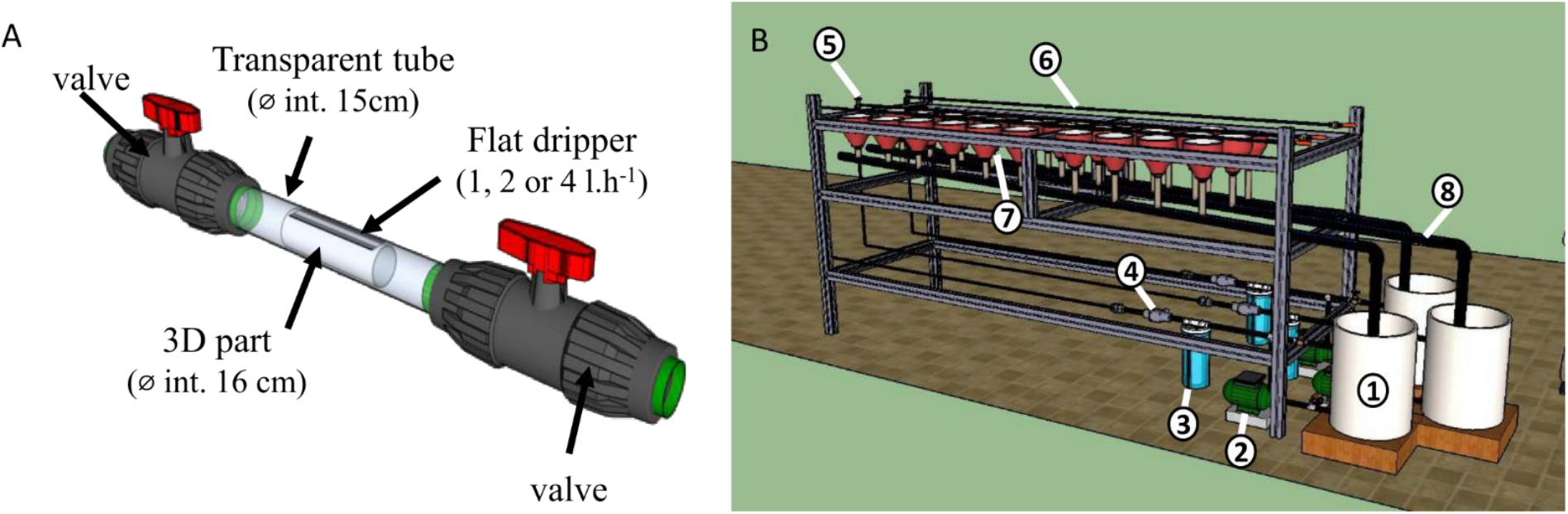
Dripper system (A) and test bench (B). The drippers were placed in a transparent tube to enable optical measurements. The test bench was composed of 1. a tank (60l); 2. a water pump; 3. a 0.13mm mesh screen filter; 4. a pressure reducer; 5. a pressure gauge; 6. the drip line with an emitter system located at 10-cm intervals; 7. a collector; 8. a gutter.

**Table 1.**
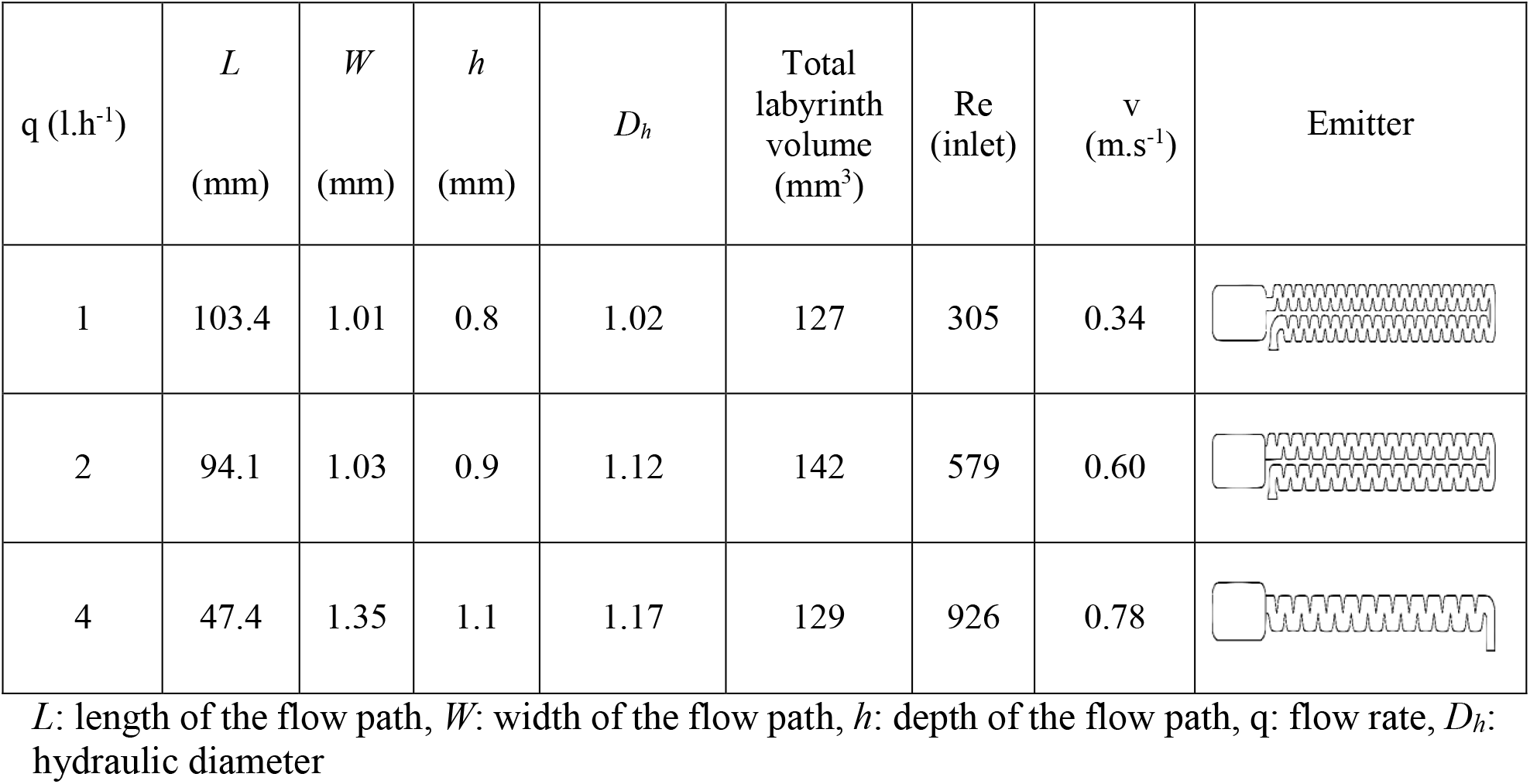
Dripper parameters

Table 1 lists the dripper characteristics. The flow regimes were characterised by a Reynolds number (Re). The Reynolds number (Re) was computed using the following formula:

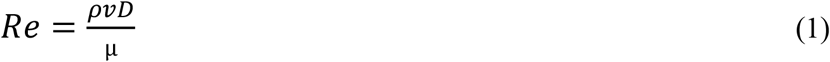

where *ρ* is water density (kg.m^-3^), *v* the water velocity across the pipe (m.s^-1^), *D* the internal diameter (m), and μ the water viscosity (Pa s).

#### 2.1.2. Computational fluid dynamic

The hydrodynamic profiles along the labyrinth channel were investigated using CFD. The numerical simulations were performed using the commercial CFD solver COMSOL Multiphysics. The geometry includes inlet and outlet channels, an outlet basin and various baffles depending on the type of dripper used (Table 1). The dimensions and channel geometries were based on the characteristics of the experimental drippers. The inlet mesh of drippers and the roughness of the walls were not included in these simulation studies. The computations were performed using a Tetrahedron grid. The influence of mesh size was tested in order to choose the appropriate computation grid. The k-ε turbulence model was used in this study. Steady state, turbulent, single-phase and incompressible fluid conditions were assumed. The fluid used was pure liquid water and was assumed not to be subject to gravity forces. Boundary conditions for both configurations were as follows: uniform pressure was imposed at the inlet (P=0.08 MPa). Atmospheric pressure was chosen for the outflow. The turbulent intensity and the hydraulic diameter were chosen for the turbulence specification method to calculate the turbulence effect (for further information on Computational Fluid Dynamics, see Pope, 2000).

#### 2.1.3. Physical-chemical and microbiological quality of the RWW

The irrigation lines were supplied with reclaimed wastewater from the Murviel-Les-Montpellier in the South of France, (43.605034° N, 3.757292° E). The wastewater treatment plant is designed around stabilisation ponds with three successive lagoons (13 680 m^3^, 4784 m^3^ and 2700 m^3^) and a nominal capacity of 1,500 Inhabitant Equivalent. The RWW was placed in a 60l tank and changed twice a week to maintain the quality close to that of the wastewater from the treatment plant. Each week (n=16), several physical-chemical and microbiological analyses were performed to evaluate the RWW quality. Chemical oxygen demand (COD), ammonia, nitrate, and phosphorus concentrations (mg l^-1^) were measured with a spectrophotometer (DR1900, Hach Company, Loveland, CO, USA) using Hach reagents^®^. Conductivity and pH were measured with probes (TetraCon^®^ 925 and pH-Electrode Sentix^®^ 940, WTW, Wilhelm, Germany). Faecal coliforms, *E. coli*, and *Enterococci* were quantified using the IDEXX method (Colilert18 and Enterolert, IDEXX Laboratories, Westbrook, ME) according to the supplier’s recommendations. The main effluent properties are listed in Table 2.

**Table 2.** Physical-chemical and microbiological characteristics of the RWW

| Characteristics | Units | Mean<br>(n=24) | SD |
| --- | --- | --- | --- |
| Soluble COD | mg l<sup>-1</sup> | 70.3 | 7.6 |
| Total suspended solids | mg l<sup>-1</sup> | 61 | 24.2 |
| Ammonia | mg l<sup>-1</sup> | 25.7 | 7.8 |
| Nitrate | mg l<sup>-1</sup> | 0.7 | 0.1 |
| Phosphorus | mg l<sup>-1</sup> | 4.9 | 0.9 |
| Conductivity | µS cm<sup>-1</sup> | 1101.4 | 51.5 |
| TS | mg l<sup>-1</sup> | 60.9 | 24.3 |
| Dissolved oxygen | mg l<sup>-1</sup> | 6.8 | 1.5 |
| pH |  | 6.9 | 0.8 |
| Total coliforms | MPN/100ml | 6.1x10<sup>5</sup> | 6.4x10<sup>5</sup> |
| *Escherichia coli* | MPN/100ml | 5.8x10<sup>4</sup> | 5.1x10<sup>4</sup> |
| *Enterococci* | MPN/100ml | 3.2x10<sup>4</sup> | 4.6 x10<sup>4</sup> |

### 2.2. Image acquisition and processing

#### 2.2.1. Image acquisition

Optical coherence tomography (OCT) was used to study the biofilm kinetics in drippers and along the milli-channel throughout the experimental period. The measurements were performed in situ and non-invasively through the transparent tube. Three-dimensional OCT measurements were acquired using a Thorlabs GANYMEDE II OCT (LSM03 lens; Thorlabs GmbH, Lübeck, Germany). OCTs have a centre wavelength of 930 nm. Due to the length of the drippers, the labyrinth channel was divided into eight parts (inlet, part 2, part 3, return, part 6, part 7, and end; Table S1 in Supplementary material) to study biofouling in different parts of the channels (Figure 2).

**Figure 2.**
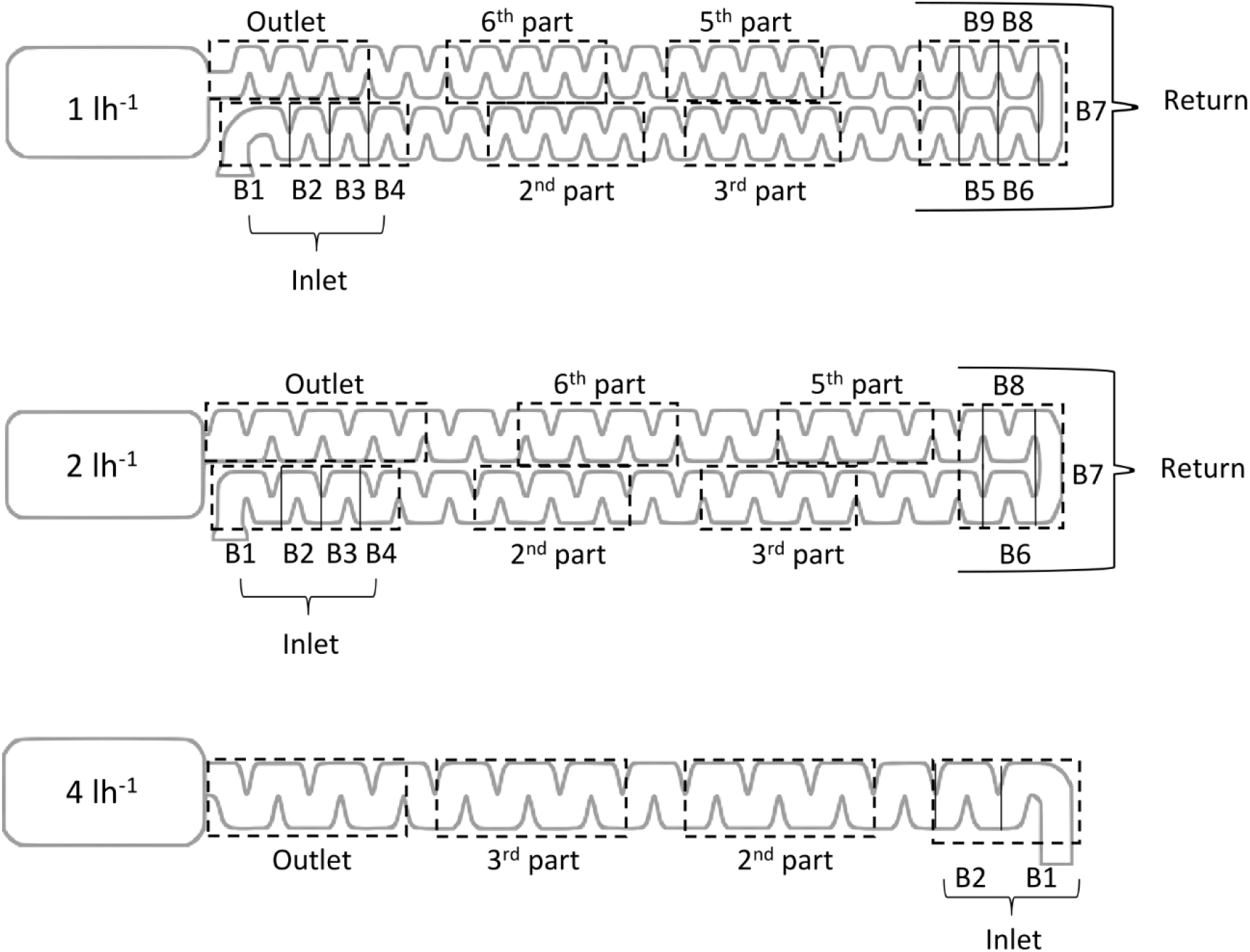
Schematic diagram of the geometric channel from the drippers. The dotted boxes correspond to the study areas. The length and width (in mm) of each baffle are listed in the table associated with Figure S1 in Supplementary material.

#### 2.2.2. Image processing

First, 3-D OCT datasets were processed in Fiji (running on ImageJ version 1.50b, Schindelin et al., (2009)) and converted into 8-bit grayscales. The datasets were resliced from top to bottom into image stacks and regions of interest (inlet and return areas) were selected. The remaining parts were allocated to the background (black). Secondly, an in-house code was used to detect the pixels associated with the plastic tube and removed using MATLAB R2018r (MathWorks ^®^, version 2018b). A threshold (adapted to each dataset) was applied to binarize the dataset and the region above the interface was quantified as biofilm. For each position (x, y), the pixels associated with the biofilm (up to the threshold) were summed (on z) to obtain the thickness of the biofilm. The biofilm volume (%) was calculated for each baffle according to Equation 2:

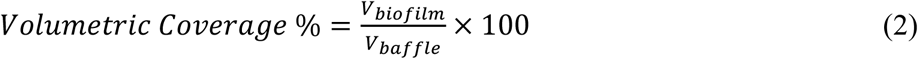

where *V_biofilm_* is the biofilm volume and *V_baffle_* is the volume of the baffle.

### 2.3. Analysis of the microbial communities

#### 2.3.1. Sampling the biofilm and reclaimed wastewater

Once a month, three drippers in each line were chosen and replaced by new ones to ensure the functioning of the system. However, these new drippers have not been analyzed. Each dripper in the transparent tube was extracted with sterile clamps, placed in a sterile tube and stored at −20°C until DNA extraction. RWW was filtered (15 to 30 mL) each week through 0.2 μm (Supor^®^ 200 PES Membrane Disc Filter, Pall Corporation) to analyse the bacterial community in the reclaimed wastewater. Filters were stored at −20°C until DNA extraction.

#### 2.3.2. DNA extraction

DNA was extracted using the PowerWater^®^ DNA Isolation Kit (Qiagen, Hilden, Germany). Samples (drippers or filters) were placed in 5 mL tubes containing beads. The manufacturer’s instructions were then followed. The DNA concentration was measured, and purity checked by spectrophotometry (Infinite NanoQuant M200, Tecan, Austria). Extracted DNA was stored at −20°C.

#### 2.3.3. Bacterial quantification by qPCR

Total bacterial quantification was performed by qPCR on biofilms from drippers targeting the V9 region from 16S rDNA. The amplification reactions were performed in triplicate, and at two dilutions to check for the absence of inhibition of the PCR reaction. Reaction mixes (12μl) contained 2.5μl of water, 6.5μl of Super Mix qPCR (Invitrogen), 100nM forward primer BAC338 (5’-ACTCCTACGGGAGGCAG-3’), 250nM of reverse primer BAC805 (5’-GACTACCAGGGTATCTAAT CC-3’) and 50nM of probe BAC516 (Yakima Yellow-TGCCA GCAGC CGCGG TAATA C –TAMRA) (Yu et al., 2005). The cycling parameters were 2 min at 95°C for pre-incubation of the DNA template, followed by 40 cycles at 95°C for 15 sec for denaturation and 60°C for 60 sec for annealing and amplification.

#### 2.3.4. Illumina sequencing

PCR amplified the V4-V5 region of 16S rRNA genes with 30 cycles (annealing temperature 65°C) using the primers 515U (5′-GTGYCAGCMGCCGCGGTA-3′) and 928U (5′-CCCCGYCAATTCMTTTRAGT-3′) (Wang and Qian, 2009). Adapters were added for multiplexing samples during the second amplification step of the sequencing. The resulting products were purified and loaded onto the Illumina MiSeq cartridge for sequencing of paired 300 bp reads according to the manufacturer’s instructions (v3 chemistry). Sequencing and library preparation was performed at the Genotoul Lifescience Network Genome and Transcriptome Core Facility in Toulouse, France (get.genotoul.fr). Mothur (version 1.39.5) (Schloss et al., 2009) was used to associate forward and reverse sequences and clustering at four different nucleotides over the length of the amplicon. Uchime (Edgar et al., 2011) was used to identify and remove chimera. Sequences that appeared less than three times in the entire data set were removed. In all, 16S rRNA sequences were aligned using SILVA SSURef NR99 version 128 (Schloss et al., 2009). Finally, sequences with 97% similarity were sorted into operational taxonomic units (OTUs) (Nguyen et al., 2016). The chloroplast sequences were removed from the raw data. Finally, BLAST (http://www.ncbi.nlm.nih.gov/BLAST/) was used to locate publicly available sequences closely related to the sequences obtained from the samples. A total of 11,357,162 reads were grouped in 7730 OTUs at the 97% similarity level.

The rarefaction curves indicated that the sequencing depths of all samples were adequate (Figure S2 in Supplementary material).

### 2.4. Statistical analyses

The kinetics of biofilm development were evaluated by modelling the biofilm formation rate using linear and spline models. The sequencing data were processed under R v3.4 (www.r-project.org) using the R-Studio (http://www.rstudio.com/) phyloseq package (McMurdie and Holmes, 2012). Kruskal-Wallis tests were performed to compare diversity and richness indices over time and between the dripper types. For the comparison of bacterial community structure, a dissimilarity matrix (Bray-Curtis) was performed and visualised using principal coordinate analysis (PCoA). A one-way analysis of similarities (ANOSIM) was used to identify significant differences in community assemblage structure between samples based on the origin of the sample (Clarke, 1993). The OTUs that contributed most to the divergence between two types of dripper were identified using Similarity Percentage (SIMPER) analysis (Clarke, 1993).

## 3. Results

### 3.1. Areas favourable for the development of biofilm

During the four month experiment, all the drippers maintained the expected flow rates, and the drippers never became clogged. In the first two weeks, the thickness of the biofilm was too low to be measured and the first measurements were made after one month. Based on OCT analysis (Figure S1, S2 and S3 in Supplemental material), the highest volume of biofilm (mm^3^) was observed in two areas: at the dripper inlet (all drippers; 2.45, 3.32 and 2.19 mm^3^ for 1, 2 and 4 l.h^-1^ drippers respectively) and in the return areas of the 1 and 2 l.h^-1^ dripper channels (with 1.62 and 2 mm^3^ of biofilm respectively, there is no return area in 4 l.h^-1^). Figure 3 shows the increase in clogging thickness in the inlet and return dripper areas at one and four months. Over time, biological fouling at the inlet area increased and gradually spread to the following baffles. The biofilm thickness in the return area increased after the large bend (B7) in both types of drippers (Figure 3C).

**Figure 3.**
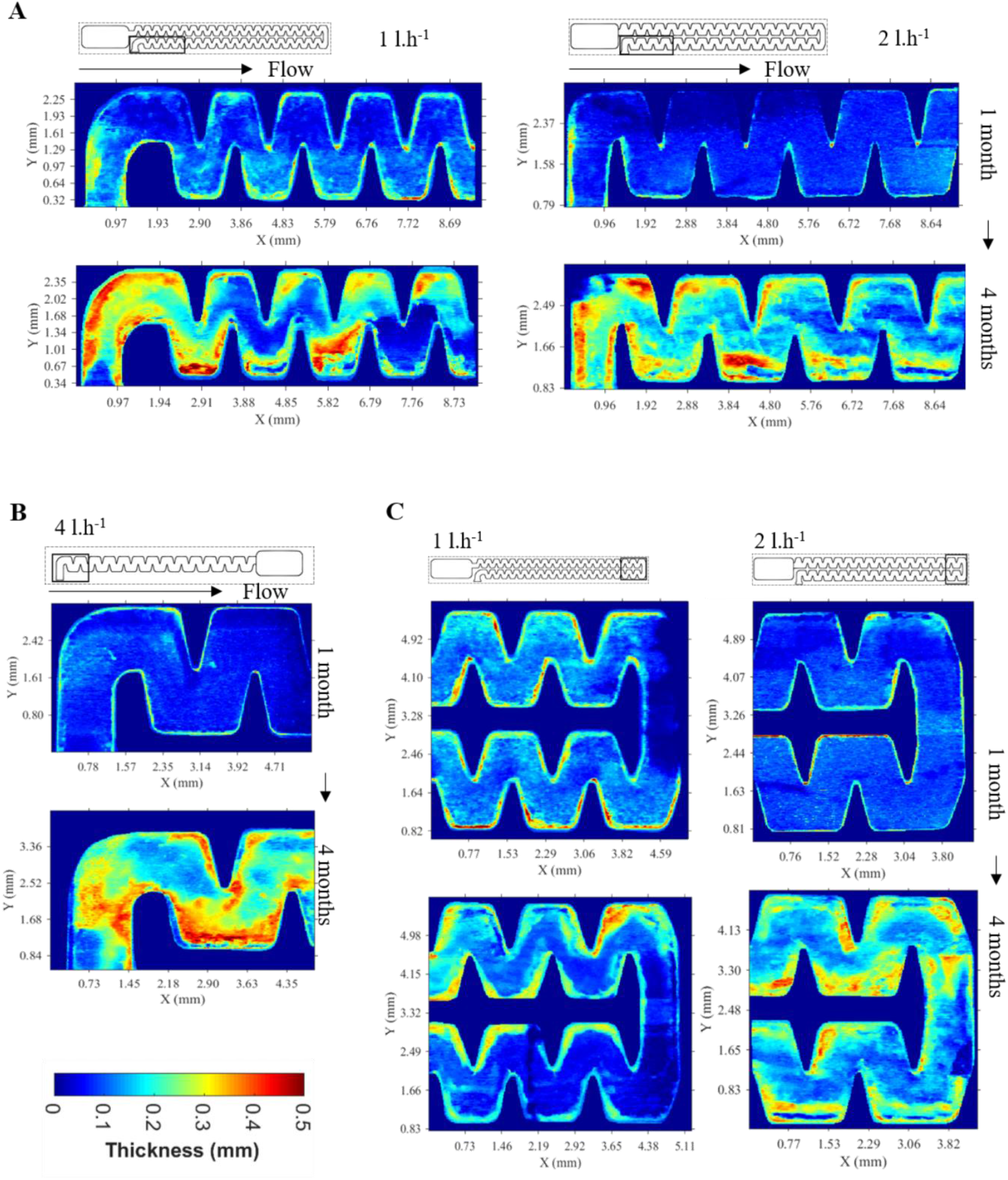
Biofilm thickness at the inlet of 1, 2 l.h-1 dripper (A) and 4 l.h-1 dripper (B) and in the return areas of drippers (C) measured after 1 and 4 months.

The mean velocity was lower at the inlet channel (Figure 4, baffle B1), as demonstrated by the CFD simulations. This low fluid velocity is the result of the rotation of the flow direction and streamline detachment, which explains why biofilm accumulated in this zone (Figure 3). Biofilm also developed in the vortex regions of the first baffles (Figure 3). According to the numerical results, low velocity (around 0.2 m.s^-1^, Figure 4) and low turbulent kinetic energy (k) values prevailed in this part of channel providing favourable conditions for the development of biofilm. Conversely, areas where k and shear stress values were higher (from the 3^rd^ baffle on) were subject to limited biofilm development.

**Figure 4.**
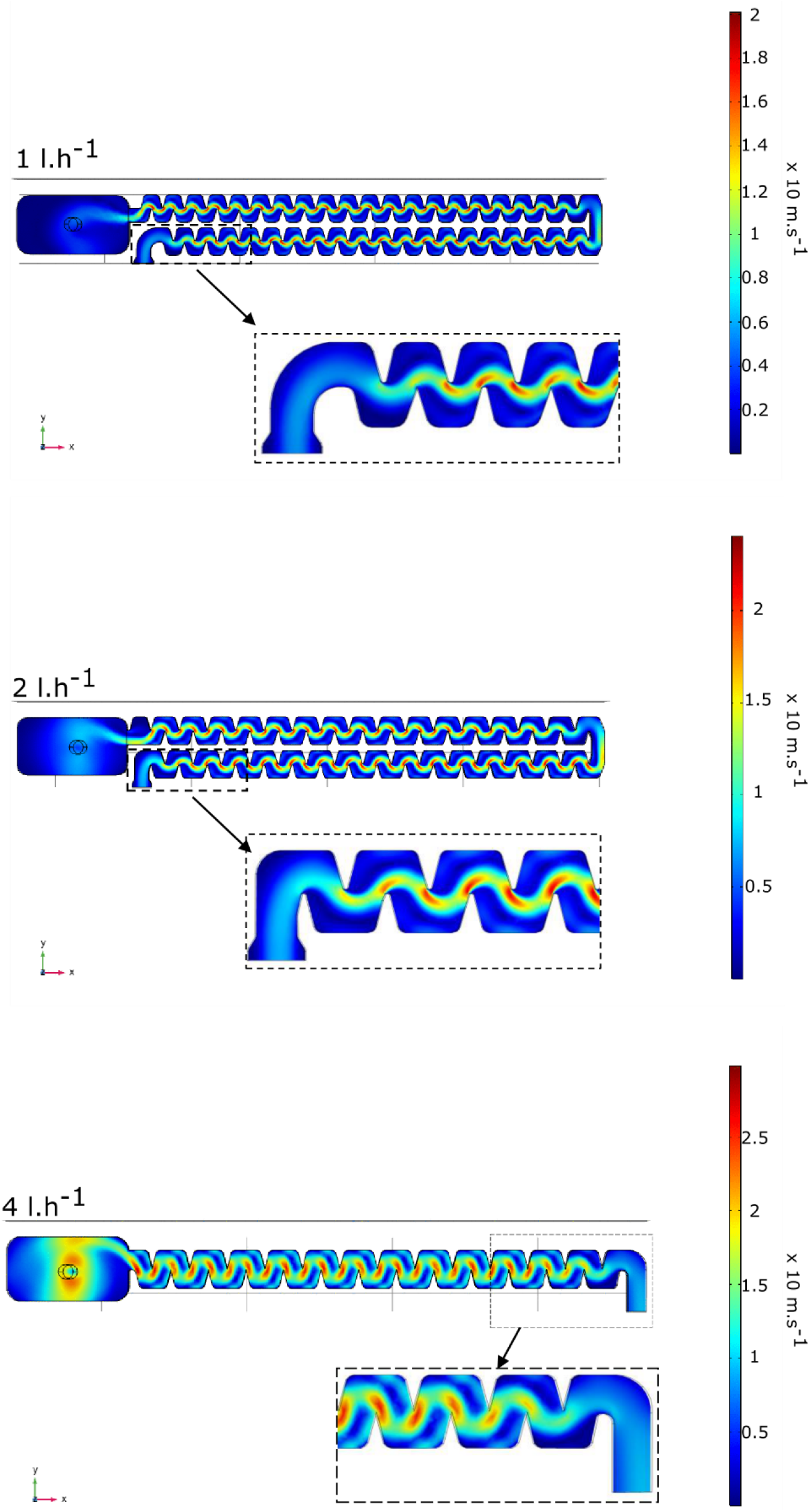
Mean velocity fields obtained numerically along the labyrinth channel at z=0 with an inlet pressure of 0.8MPa

The volume of the baffles differed with the dripper. Therefore, to compare the level of clogging between drippers, the biofouling volume was normalized by the volume of the corresponding baffle (Equation 2). Mean clogging of the inlet baffles increased over time in all three types of drippers (Figure 5). The level of biofouling were significantly influenced by the type of dripper (Chi^2^ = 115, p-value < 0.05) and the time (Chi^2^ = 54, p-value < 0.05). The level of biofouling increased more rapidly in 1 and 2 l.h^-1^ drippers than in 4 l.h^-1^ drippers (Wald test, p-value < 0.05).

**Figure 5.**
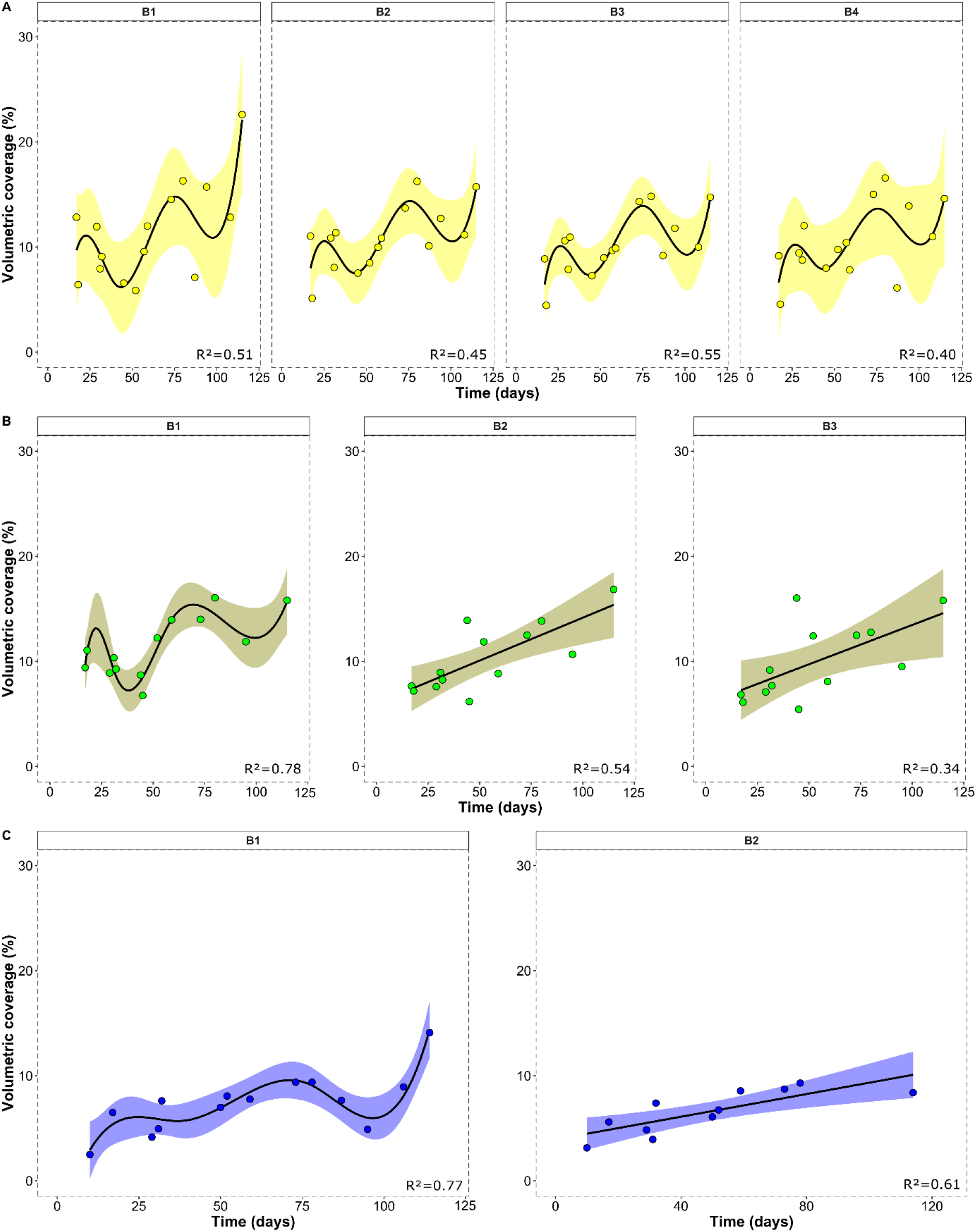
Mean volumetric biofilm coverage at the inlet of the drippers 1 l.h^-1^ (A) and 2 l.h^-1^ (B) and 4 l.h^-1^ (C) over time. The lines and confidence interval (95% of associated variance) correspond to the spline and linear model used for each dripper type (Baffle B4 from 2 l.h^-1^ was not included due to the too low regression value (R^2^<0.25).

There was no statistical effect of baffles on volumetric coverage over time. However, the volume of biofouling in the first baffle was significantly higher in 1 l.h^-1^ drippers than in 2 l.h^-1^ and 4 l.h^-1^ drippers after four months with a mean volumetric coverage of 22.6%, 15.8% and 14.1 % for 1 l.h^-1^, 2 l.h^-1^ and 4 l.h^-1^ drippers, respectively (Kruskal test, p-value < 0.05). The mean increase in biofouling in the 1 l.h^-1^ drippers presented a sinusoidal dynamic with up and down phases at the inlet baffle. This could mean several detachment events occurred. The same cycle was observed in the 2 and 4 l.h^-1^ but only in the first baffle. The detachment event occurred at ~ 3 months in the 4 l.h^-1^ drippers whereas the first event appeared at 1 month in the 2 l.h^-1^ drippers. After this baffle, the increase in biofouling was linear. At the return zone in 1 and 2 l.h^-1^ drippers, the volumetric coverage increased over time but was heterogeneous. Thus it was not possible to define kinetic models.

The type of dripper (flow rate and flow cross-section) influenced the level of clogging of the labyrinth with a higher clogging rate in 1 l.h^-1^ drippers (lower flow rate and narrower flow cross-section).

### 3.2. Microbial diversity differed between dripper biofilms and reclaimed wastewater

The structure of the bacterial communities in biofilms collected in the three types of drippers were compared using 16S rDNA Illumina sequencing. The bacterial communities in the biofilms were also compared with the communities present in reclaimed wastewater. The chloroplast sequences were removed from the raw data and represented 1.6% and 5.4% of the total sequences in biofilms and in RWW samples, respectively. Microscopic observations of the biofilms in the pipes (data not shown) also revealed the presence of eukaryotic microorganisms (bacterial predators) but the biofilms were still mainly composed of bacteria.

The biofouling of the drippers increased over time in all three types of drippers as shown by the increase in DNA concentration. (Table 3). On the other hand, the number of bacteria remained unchanged over time in the three types of drippers and already reached 10^9^ copies of 16S rDNA/dripper at 32 days in all three types of dripper.

**Table 3.** DNA concentration and quantification of bacteria by qPCR in dripper biofilms

| Sampling time (days) | Dripper flow rate (l.h<sup>-1</sup>) | Mean DNA concentration (µg/µl) | Mean n° of bacteria /dripper (16S copies/dripper) |
| --- | --- | --- | --- |
| 32 | 1 | 137.4±67.9<sup>a</sup> | 6.2±0.9 x 10<sup>9a</sup> |
|  | 2 | 113.9±42.7<sup>a</sup> | 5.7±0.6 x 10<sup>9a</sup> |
|  | 4 | 118.1±26.8<sup>a</sup> | 9.9±1.2 x 10<sup>9a</sup> |
| 72 | 1 | 220.4±34.6<sup>a</sup> | 9.2±0.5 x 10<sup>9a</sup> |
|  | 2 | 287.4±113.8<sup>a</sup> | 1.2±0.05 x 10<sup>10a</sup> |
|  | 4 | 194.4±20.2<sup>a</sup> | 4.3±2.3 x 10<sup>9a</sup> |
| 115 | 1 | 325.2±70.8<sup>a</sup> | 8.7±0.4 x 10<sup>9a</sup> |
|  | 2 | 387.8±164.1<sup>a</sup> | 1.4±0.5 x 10<sup>10a</sup> |
|  | 4 | 346.1±34.5<sup>a</sup> | 5.4±1.4 x 10<sup>9a</sup> |

Kruskal-test and Wilcoxon ad hoc tests were performed on each sampling occasion to compare the values between the drippers (n= 3 for each type of dripper on each sampling event); letters show Wilcoxon test results.

The richness and diversity indices of the biofilms did not significantly differ between the three types of drippers, but increased over time in all three (Table 6). The richness and diversity indices of RWW were lower than those in the dripper biofilms (Table 4). The majority of operational taxonomic units (OTUs) were shared by all the biofilms (Figure S5 in Supplemental material). Some OTUs and bacterial genera were specific to the dripper flow, but their relative abundance was less than 0.1% (Table S2 in Supplementary material).

**Table 4.**
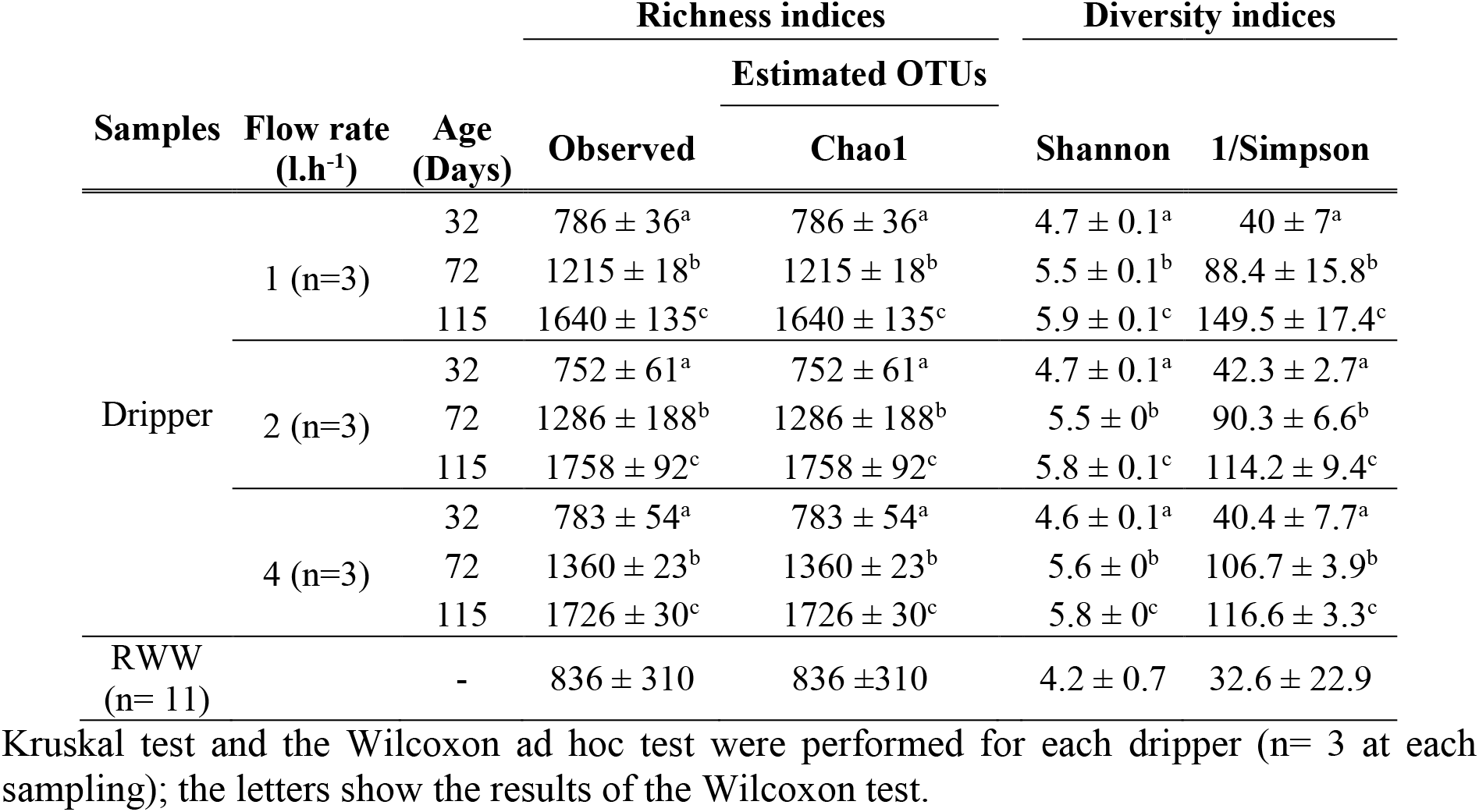
Estimated species richness and diversity indices according to the dripper flow rate and the RWW

Proteobacteria, Bacteroidetes, Firmicutes and Chloroflexi were the main phyla in the dripper biofilms (Figure 6A). The main phyla found in RWW were Proteobacteria (25-55%) mainly composed of β-Proteobacteria (10-34%) and γ-Proteobacteria (2-34%), followed by Bacteroidetes (22-40%) and Actinobacteria (7-14%) (Figure 6B). These differences indicate that among bacteria in the reclaimed wastewater, bacteria able to attach to the surface of the dripper under the corresponding flow regime were selected, especially among Firmicutes and Chloroflexi (Figure 7). From 49 days onwards, cyanobacteria phylum appeared in reclaimed wastewater, explaining the increase in cyanobacteria in the drippers on the second sampling occasion (72 days). The same observation was made concerning Spirochaetae and Chlorobi phyla. Thus, RWW influenced the bacterial composition of the dripper biofilms. The structure of the biofilms in the drippers evolved over time according to changes in the microbial diversity in the reclaimed wastewater, and with the selection of some adapted taxa able to adhere to the surfaces, such as filamentous Chloroflexi bacteria.

**Figure 6.**
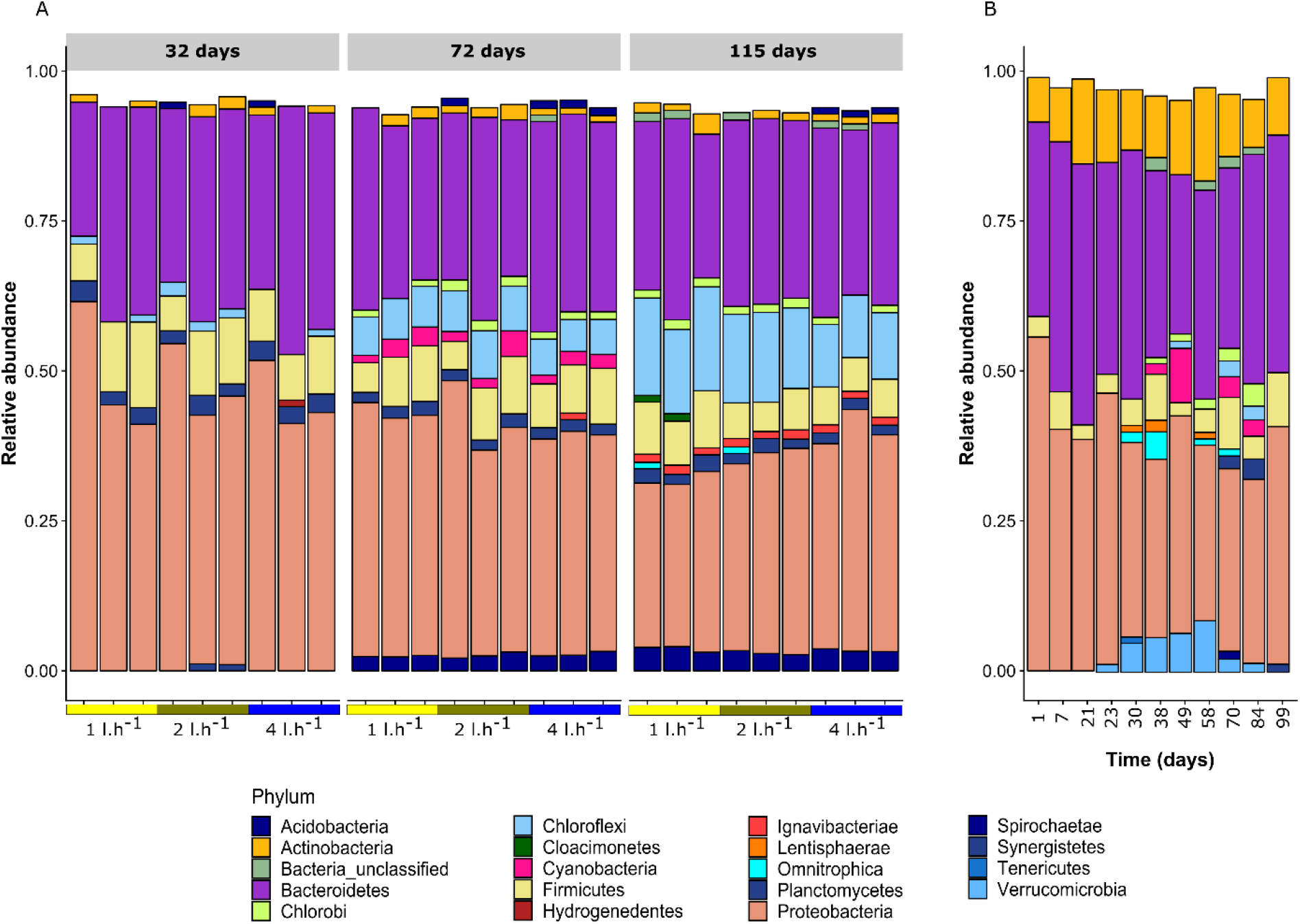
Relative abundance of bacterial phyla (>1%) in dripper biofilms (A) and in reclaimed wastewater (B).

### 3.3. The dripper flow parameters influenced the composition of the bacterial communities

The impact of flow parameters on the structure of bacterial communities was first investigated at the phyla and familly taxonomic levels. Proteobacteria phylum was the most abundant phylum in the biofilms (Figure 6) and was mainly composed of α-Proteobacteria, β-Proteobacteria (mainly composed of members of the Comamonadaceae family), γ-Proteobacteria and δ-Proteobacteria. At 32 days, the mean abundance of Proteobacteria was similar in all dripper types, with 49, 47 and 45% in the 1, 2 and 4 l.h^-1^ dripper biofilms, respectively (Kruskal-test, p-value > 0.05). At 115 days, the mean relative abundance of Proteobacteria (and more specifically β-Proteobacteria) was significantly lower in the 1 l.h^-1^ dripper biofilms than in the 2 and 4 l.h^-1^ dripper biofilms (Kruskal-test, p-value < 0.05) and decreased significantly until 28% vs 33 and 37% for the 2 and 4 l.h^-1^ dripper biofilms, respectively.

The phylum of Bacteroidetes was mainly composed of Sphingobacteriia (Chitinophagaceae, Lentimicrobiaceae, Saprospiraceae and Sphingobacteriaceae class members), Bacteroidia (mainly Draconibacteriaceae and Rikenellaceae member families) and Bacteroidetes_vadinHA17. There was no effect of the type of dripper on the mean relative abundance of Bacteroidetes (~ 30% over time in each dripper type, Kruskal test, p-value > 0.05). Over time, the relative abundance of Sphingobacteriia and Bacteroidia decreased while the relative abundance of Bacteroidetes_vadinHA17 increased in the three types of drippers.

The mean relative abundance of Chloroflexi (mainly composed of Anaerolineaceae and Caldilineaceae member family) increased significantly over time and was significantly higher in 1 l.h^-1^ dripper biofilms after 115 days (16% versus 14% and 10% for 1, 2 and 4 l.h^-1^ dripper biofilms, respectively, Kruskal test, p-value < 0.05). At 115 days, the mean relative abundance of the Anaerolineaceae family was significantly higher in 1 l.h^-1^ dripper biofilms than in the 4 l.h^-1^ dripper biofilms (Kruskal test, p-value < 0.05). There were no significant differences in the Firmicutes phylum (mainly represented by Christensenellaceae, Clostridiaceae and Ruminococcaceae), according to the dripper types or over time (Kruskal test, p-value > 0.05). The impact of flow rates on bacterial communities was then investigated at the genus level. Table 5 lists the 10 top genera found in all the dripper biofilms over time.

**Table 5.**
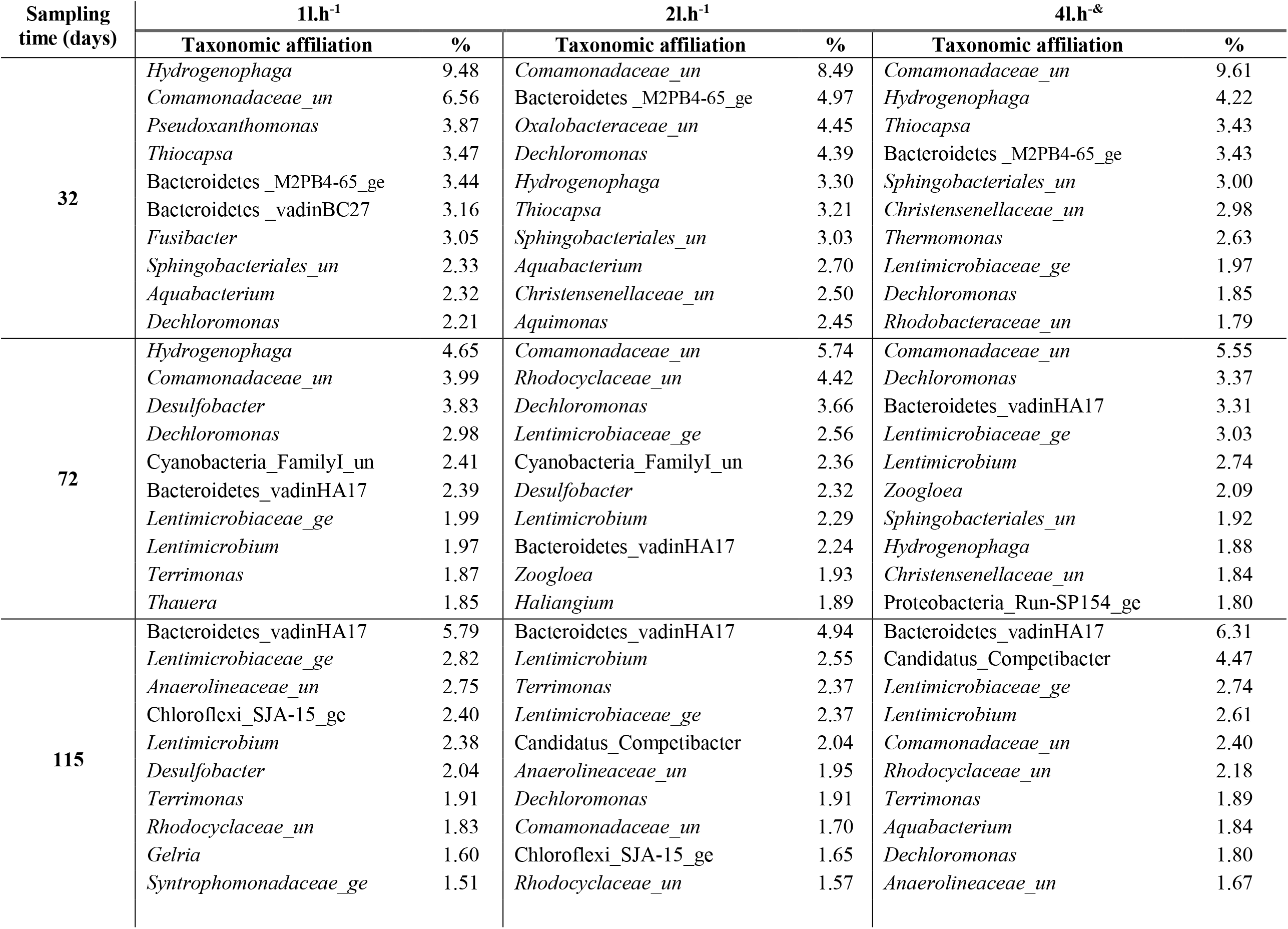
Ten most abundant genera found in dripper biofilms according to the sampling time. ‘_un’: For unclassified genus (‘_un’), the taxonomic affiliation is given at family level _ge: uncultured bacteria identified at the genus level by molecular studies.

At 32 days, the 1 l.h^-1^ dripper biofilms were dominated by the Comamonadaceae family (6%) including *Hydrogenophaga* genus (9%) and *Pseudoxanthomonas* (3%). The mean relative abundance of the *Hydrogenophaga* genus was significantly influenced by the flow rate (Kruskal test, p-value < 0.05) and was higher in the 1 l.h^-1^ dripper biofilms (3% and 4% for 2 and 4 l.h^-1^ dripper biofilm, respectively). Over time, the mean relative abundance of members of the Comamonadaceae family and of the *Hydrogenophaga* genus decreased significantly in the 1 l.h^-1^ (Kruskal test, p-value < 0.05), close to the relative abundance in 2 and 4 l.h^-1^ dripper biofilms at the end (Kruskal test, p-value > 0.05). The relative abundance of Bacteroidetes_vadinHA17 and members of the Lentimicrobiaceae family, which belong to Bacteroidetes, increased significantly over time in each dripper type (Kruskal test, p-value < 0.05) and dominated in all three drippers at 115 days (Kruskal test, p-value > 0.05).

Even if the majority of the genera were the same in the different drippers and on the sampling occasions, some genera with low abundance (<1%) depended specifically on the dripper flow rate and the sampling time (Table 8, Supplementary Figure 9). For instance, only *Vulcaniibacterium* and *Microvirga* genera were found in the 1 l.h^-1^ dripper on each sampling time. This means that hydrodynamic conditions influenced both the dominant and less abundant bacterial taxa.

### 3.4. Dripper hydraulic properties impact bacterial community structure

PCoA was performed to compare the structure of bacterial communities between the three types of dripper over time and confirmed that the bacterial community in the dripper biofilms changed depending on the type of dripper and over time (Figure 7). At 31 days, the bacterial populations in 2 and 4 l.h^-1^ were clustered (one-way ANOSIM, R=0.71, p < 0.05). The bacterial communities in the 1 l.h^-1^ dripper biofilms formed a second cluster (Figure 7, A). This means that the hydraulic conditions had little influence on the structure of the microbial community in 2 and 4 l.h^-1^ compared to that in the 1 l.h^-1^ dripper biofilms at the first stage of development. At the end of the experiment (Figure 7, C) the bacterial community in 2 and 4 l.h^-1^ diverged and formed specific groups (one-way ANOSIM, R=0.65, p<0.05).

**Figure 7.**
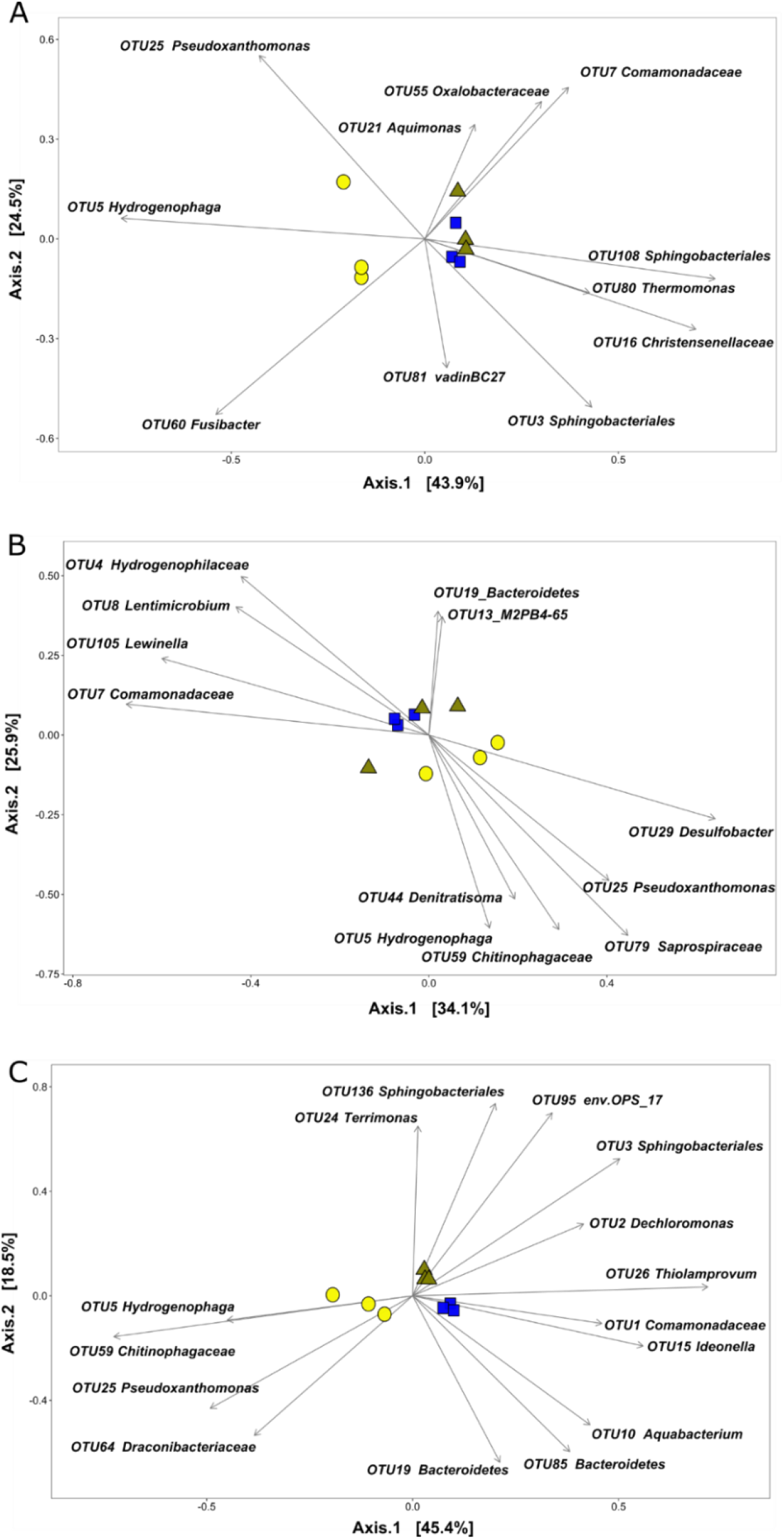
PCoA ordination plot of bacterial communities found in 1 l.h-1 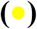, 2 l.h-1 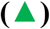 and 4 l.h-1 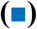 dripper biofilms at 32 days (A), 72 days (B) and 115 days (C). The analysis was based on Bray-Curtis similarity coefficients. Arrows indicate the orientation and contribution of the OTUs (most significant contributors explaining 50% of the global bacterial divergence).

The main contributors (those explaining 50% of the global bacterial divergence are indicated in Figure 7) of the bacterial community were identified using SIMPER analyses and were shown to change over time. Those explaining 50% of the global bacterial divergence are indicated in Figure 7. At 32 days, 11 OTUs played a significant role in bacterial community divergence. *Pseudoxanthomonas* (OTU 25), *Hydrogenophaga* (OTU 5) and *Fusibacter* genera (OTU 60) were the main contributors and were associated with the 1l.h^-1^ drippers. These genera were also the main genera found in 1 l.h^-1^ dripper biofilms at 32 days (Table 4). At 115 days, 23 OTUs significantly contributed to divergence. Hydrogenophaga (OTU 5) and *Pseudoxanthomonas* (OTU 25) were still strong contributors associated with 1 l.h-1 dripper biofilms. *Aquabacterium* (OTU 10), *Ideonella* (OTU 15) and Bacteroidetes members (OTU 19, OTU 85) were associated with the 4l.h^-1^ dripper biofilms, whereas Sphingobacteriales family members (OTU 3, OTU 136) and *Terrimonas* genus (OTU 24) were associated with 2l.h^-1^ dripper biofilms.

## 4. Discussion

The main problem with the use of the drip irrigation supplied with reclaimed wastewater is biofouling. Understanding the development of biofilms is the only way to limit the clogging of drip irrigation systems. In this study, a laboratory experiment was performed to investigate the kinetics of biological clogging and changes in the bacterial communities over time as a function of the hydrodynamic parameters (flow rate, cross-section) of the drippers. Three types of commercial flat drippers used in agriculture with specific channel geometry and different flow rates (1, 2 and 4 l.h^-1^) were installed on a test bench and supplied with treated urban wastewater.

Based on OCT results, the thickness and the volume of biofilm were higher in the inlet of the channel (mostly in the first baffle) in all the drippers and close to the return areas of the labyrinths in 1 and 2 l.h^-1^ drippers (Figure 3 and Figure 5). Simulation of the flow of the milli-channel showed that the fluid velocity and kinetic turbulence energy were lower in these areas (Figure 4). This is consistent with the results obtained by Liu et al., (2019) who analysed the effect of water velocity and nutrient concentration on the development of biofilms in a microchannel device. These authors showed that an increase in velocity from 1.66 m.s^-1^ to 4.17 m.s^-1^ gradually decreased the coverable biofilm over time, increased the nutrient uptake to the biofilm and created a thinner biofilm. The reduction in the velocity at the inlet and return areas can facilitate adhesion and deposition on the surface of the wall (Ait-Mouheb et al., 2018), explaining the sensitivity to clogging of these areas. High shear stress and high-water velocity thus reduce the biofilm maturation process. Biofouling was also more abundant at the corners of the milli-channel from the inlet (Figure 3), where the water velocities and shear stress were lower (Figure 4). It is interesting to note that biofouling also decreased in the vortex zones along the channel even though there was no change in flow behaviours. Flow velocity can influenced the mass transfer (nutrients) and shear stress (Araújo et al., 2016; Liu et al., 2019) along the channel. Monitoring the diffusion of nutrients to the biofilm along the milli-channel from drippers fed with RWW would improve our understanding of the formation and development of clogging.

The biofouling of the dripper increased over time in all three dripper types tested here but the 1 l.h^-1^ drippers were more sensitive than the 2 and 4 l.h^-1^ drippers (Figure 5). The cross-section of 1 l.h^-1^ drippers (1.02 mm^2^) is smaller than that of the two others (2 l.h^-1^= 1.12 mm^2^ and 4 l.h^-1^ = 1.17 mm^2^). Previous studies have shown that a high flow rate and large cross-section reduced clogging of the drippers (Duran-Ros et al., 2009; Ravina et al., 1992). Qian et al. (2017) investigated the development of biofouling of a milli-channel system fed with different quality treated wastewater (secondary, tertiary) and observed that the global growth of biofouling increased linearly over time. In the present study, clogging was monitored by OCT in each baffle of the inlet channel. Thus, we saw for the first time that dripper clogging can alternate between up and down phases, mainly in the first inlet baffles. The down phases may be linked with detachment events (Flemming and Wingender, 2010).

Detachment events appeared to occur more frequently in the 1 l.h^-1^ drippers (Figure 5). The flow characteristic (water velocity, shear stress) could explain the detachment events. Araújo et al., (2016) noted that *Pseudomonas fluorescens* species increased the production of extracellular polymeric substances (EPS) with an increase in water flow velocity (0.1 m.s^-1^ to 0.8 m.s^-1^), which resulted in thinner and denser biofilms. EPS are essential to ensure the maintenance and stability of the biofilm on a surface. Biofilms grown under low velocities are subject to lower shear stress forces which favour faster growth (Melo and Vieira, 1999). However, biofilms subjected to low flow constraints have limited mechanical strength and are more prone to sloughing than those formed at higher flows (Teodósio et al., 2011). Even if the regime was turbulent in all three types of drippers studied here, differences in the Reynolds number and in the water velocities at the channel inlet can influence EPS production and may explain the differences in detachment events observed between the drippers. Biological processes including effect of bacterial predators can also cause detachment of the biofilm (Dawn Parry, 2004). Further research on the effect of the internal forces on the biofilm structure (density, porosity), EPS production and microbial composition will advance our understanding of biofilm growth phenomena in drippers.

The structure of bacterial community of the RWW has a direct influence on the composition of the biofilms. Members of the Comamonadaceae family (β-Proteobacteria including *Hydrogenophaga* genus) dominated in the biofilm samples taken at 32 and 72 days, but were replaced by the Bacteroidetes_VadinHA17 wastewater sludge group belonging to Bacteroidetes at the end (Table 5). This non-motile anaerobic Gram-negative group is often found in sludge and indicated that over time, bacteria commonly found in wastewater settled and became the majority in communities. Comamonadaceae member family, which includes many denitrifier members, were also commonly found in wastewater treatment plants and were involved in the first phases of clogging of membrane bioreactors fed with wastewater (Ziegler et al., 2016). Other phyla such as Spirochaetae and Cholorobi appeared in the RWW over time and eventually in the biofilms (Figure 6). The RWW used in this study was obtained after treatment by lagooning and the treatment process can affect the structure and bacterial composition of biofilms. Further research is then needed to study the effect of the differentwastewater treatments on biofilm composition in irrigation lines in order to improve management of biofilm development in the drippers.

Diversity and richness did not influenced by hydraulic parameters of the drippers. This is not consistent with Shi et al. (2019) who studied the effect of hydraulic regime on biofilm evolved in pipes from drinking water distribution systems. However, these authors used a wider range of flow rates (120 to 600 l.h^-1^). The differences between the drippers in terms of flow rate may not be sufficient to observe an effect on these indices.

Although bacterial diversity was not influenced by the hydraulic parameters, the structure of the bacterial community was. This is in agreement with the results of previous studies (Ai et al., 2016; He et al., 2019; Saur et al., 2017). The level of clogging decreased with an increase in water velocity (0.34, 0.61 and 0.78 m.s^-1^ for 1, 2 and 4 l.h^-1^ drippers respectively) and shear stress in the channel. However the structure of bacterial communities appeared to be less affected by water velocity above 0.61 m.s^-1^ (2 l.h^-1^ dripper). Even though previous studies have shown that the increase in velocity and shear forces influence the spatial organization and structure of the bacterial community (Saur et al., 2017), the differences between the 2 and 4 l.h^-1^ drippers may not be sufficient to influence this community in the first phases of biofilm development. This could explain the similarities in the structure of the bacterial communities sampled in 2 and 4 l.h^-1^ drippers compared to the 1 l.h^-1^ dripper biofilms until 115 days.

Proteobacteria, Bacteroidetes, Firmicutes and Chloroflexi were the most abundant phyla in all three types of dripper (Figure 6). These phyla have been already been described in drippers supplied by treated wastewater (Lequette et al., 2019; Song et al., 2019; Zhou et al., 2013) and play a key role in biofilm development. Over time, the relative abundance of Proteobacteria and more specifically β-Proteobacteria decreased significantly in 1 l.h^-1^ dripper biofilms compared to 2 and 4 l.h^-1^ dripper biofilms. β-Proteobacteria have been reported to be a dominant group in both drinking water biofilms (Douterelo et al., 2013) and wastewater biofilms (Biswas and Turner, 2012; Ma et al., 2013) and could play an important role in the formation of biofilm in drippers. This group can easily attach to surfaces and can withstand high water velocity and high shear force (Douterelo et al., 2013), which may explain their dominance in drippers 2 and 4 l.h^-1^. Chloroflexi and Bacteroidetes phyla were also in the majority in dripper biofilms. These phyla are commonly found in activated sludge of wastewater treatment plants and include filamentous bacteria as Anaerolineaceae family members (Chloroflexi) or Saprospiraceae family members (Bacteroidetes).

Research has shown that filamentous bacteria are essential for the formation of activated sludge but that, in too high a concentration, they are responsible for foaming and bulking events (Kragelund et al., 2008, 2007; Nielsen et al., 2009) and for clogging membrane bioreactors in wastewater treatment plants (Li et al., 2008). Thus, we showed for the first time that filamentous bacteria have a key role in the biofouling of the drippers. The relative abundance of filamentous bacteria was also influenced by hydraulic properties as the flow rate of the drippers. Anaerolineaceae family members (Chloroflexi) were more abundant in 1 l.h^-1^ dripper biofilms at the end than in 2 and 4 l.h^-1^ drippers and could have a key role in biofouling of drippers.

The increase of Chloroflexi over time can also be explained by the interaction with other bacteria. Chloroflexi and *Hydrogenophaga* spp (Proteobacteria phylum), mainly found in 1 l.h^-1^ drippers, were also observed by Ziegler et al. (2016) who studied the biofouling of a pilot-scale membrane bioreactor fed with wastewater. From a physiological point of view, *Hydrogenophaga* spp. are able to synthesise polymeric substances (Calderer et al., 2014). Filamentous Chloroflexi produces a complex of enzymes able to degrade EPS, in turn, enabling the cells to use EPS as substrate to maintain their activities (Kragelund et al., 2007). EPS produced by *Hydrogenophaga* spp could facilitate fouling due to filamentous bacteria and flow parameters. Thus, the increase in Chloroflexi abundance over time could be explained by a sufficient EPS rate and by the presence of *Hydrogenophaga* spp at the early stage biofilm formation.

The divergence between the 1 l.h^-1^ drippers and the 2 and 4 l.h^-1^ drippers prevailed over time and was mainly driven by *Hydrogenophaga* and *Pseudoxanthomonas* genera (Figure 7). Although no clear information is available on the effect of hydraulic conditions on the installation of these two bacterial genera, they are often found in biofilms associated with membrane reactor clogging supplied by wastewater (Zheng et al., 2018; Ziegler et al., 2016). Contrary to the 1 l.h^-1^ dripper biofilms, biofilms from the 4 l.h^-1^ dripper were mainly driven by *Aquabacterium* (Proteobacteria) and *Ideonella* (Proteobacteria, Comamonadaceae family) genera after 4 months. These genera were already found associated to wastewater biofilms (Lequette et al., 2019; Luo et al., 2017; McCormick et al., 2016) and included species able to metabolize plasticizers used in plastics (Kalmbach et al., 1999; Tanasupawat et al., 2016). Thus the type of material and the hydraulic parameters of the drippers could influence the presence of bacteria with specific metabolic activities over time.

## 5. Conclusion

The combined use of OCT and high throughput sequencing highlighted the influence of hydrodynamic parameters (flow rate, cross-section) of the drippers supplied with RWW on biofilm development and bacterial communities:

1. Biofouling was facilitated close to the inlet and in the vortex zones of the dripper channels, more specifically in 1l.h^-1^ drippers, due to lower turbulence kinetic energy and water velocity.
2. The water velocity influenced the structure of bacterial communities in biofilms. However, the slight differences between 2 and 4 l.h^-1^ would explain the similarities in the structure of the bacterial community compared to those of the 1 l.h^-1^ dripper biofilms.
3. The low flow velocities in the 1 l.h^-1^ drippers seemed to favour the presence of *Hydrogenophaga* and *Pseudoxanthomonas* genera in the first stage of biofilm development and the installation of filamentous bacteria belonging to Chloroflexi phylum at the end.

Water velocity evolves along the channel with low velocities at the inlet and in the vortex zones. These changes can influence the transport of nutrients to the biofilm along the channel. Further studies on nutrient transport and on the structure of the bacterial community of the biofilms along the drippers should increase our understanding of clogging phenomena in sensitive areas.

## Nomenclature

OCT: Optical coherence tomography
RWW: Reclaimed wastewater
s-COD: soluble Chemical Oxygen Demand
CFD: Computational fluid dynamic
TS: Total solid

## Acknowledgements

The authors gratefully acknowledge the financial support of the French Water Agency, project “Experimental platform for the reuse of reclaimed wastewater in irrigation, Murviel-lès-Montpellier”. We thank Guillaume Guizard (LBE-INRA, Narbonne, France) and Jean-François Bonicel (IRSTEA, Montpellier, France) for their contribution to the development of the dripper system, Annabelle Mange (IRSTEA, Montpellier, France) and Valérie Bru-Adam (LBE-INRA, Narbonne, France) for their assistance in the field and at laboratory.

